# Mechanical unfold and transport of Green Florescent Protein through a nanopore

**DOI:** 10.1101/294819

**Authors:** Muhammad Adnan Shahzad

## Abstract

We report the unfold and trans-location of Green Fluorescent protein (GFP) mechanically by a constant force acting parallel along the axis of nanopore. A coarse-grained numerical model (Go-model) were implemented both for the protein and the nanopore. Detail description of each peptide unfold by the constant force is presented. Depending on the GFP topological structure, *β*-sheet barrel, the protein unfold and transport as a double loop conformation in the confinement geometry. The result is compared with maltose binding protein (MBP), having majority of alpha helix, which unfold and trans-locate as single profile conformation through nanopore. The result emphasis that protein with different topological structure unfold and trans-locate in different fashion depending on their native fold structure.

## I. INTRODUCTION

Recently, many efforts has been made to comprehend the unfolding of bio-molecules using mechanical force. Atomic force microscopy (AFM) or optical tweezers methods are implemented to study experimentally the unfolding trajectories of proteins [1–5]. The proteins are stretched in atomic force microscopy in order to characterized folding force. Many efforts has been made to understand equilibrium and non-equilibrium behavior of proteins using single molecule method and molecular dynamics simulation. Single molecule methods is also employed to study the unfolding and refolding of protein inside a nanopore in electric field [6–10, 12–18]. The nanopore technique allows one to probe the conformational space of proteins. Several studies with protein channel (*α*-hemolysin) or solid-state nanopores have been able to detect the unfolding process with some experiments even generating unfolding curves of wild type and/or mutant proteins using event frequency analysis. Single molecule mechanical unfolding studies have shown that mechanical extended through the N- or C- termini, proteins reveals a wide variety of mechanical behavior. *β*-sheet protein show more mechanically resistant than their *α*-helical counterparts. We present through molecular dynamics simulation using coarse grained model that the beta-barrel protein (GFP) possessed more mechanical resistance during the unfold and trans-location process through a nanopore.

Green fluorescent protein (GFP) [20] is among one the most interesting protein studies experimentally with force spectroscopy technique. Green fluorescent protein (GFP) from the jellyfish Aequorea victoria is currently used in biological and medical research as a marker of gene expression and protein localization, as an indicator of protein-protein interactions and as a bio-sensor [21]. The GFP contains 238 residues with 11-stranded *β*-barrel structure. Many experimental research group has reported the mechanical properties of proteins.

In [23], the experimental results of the group of Oukhaled. et. al. shows that a partially folded protein trans-locating through nanpore exhibit very long block-ades current in nanopore. Motivated by such experiment and the mechanical behavior of *β*-sheet and α-helical proteins, here we report the unfold and trans-location of GFP (*β*-barrel protein) through nanopore using coarse gain molecular dynamic simulation in comparison with maltose binding protein. The GFP initially unfold completely up to *β*6-beta sheet and than enter as partially folded protein in nanapore. The partially folded protein transport through nanppore as a loop conformation. The trans-location of partial folded GFP exhibit a pause or rapture effect in the trajectory of protein unfolding. Experimentally, this behavior is treated as long blockages current when trans-locating a particaly folded protein through a biological or solid state nanopore.

The paper is organized as follows: sec.II we derscibe the computational model used to simulate GFP unfolding in a confinement cylindrical geometry. We used coarse grained Go model both the protein and confinement geometry. In sec.III we discuss the results of mechanical unfolding of GFP in comparison with Maltose-binding protein consists of majority of *α*-helix. Sec. IV is devoted to the conclusion.

## II. MODEL AND SIMULATION DETAILS

### A. Protein and nanopore model

We used a Gō-like force field acting on the beads in our numerical simulation which preserve the secondary structure content, the beta sheets and alpha helices, of a protein chain [22, 24, 25]. In this approach we used the following potential acting on the material point of the protein: (1) Peptide potential (or bond potential) *V*_*p*_, (2) Bending angel poteila *V*_*θ*_, (3) Twisted angel potential *V*_*φ*_ and (4) Non-boned interaction (Lennard-Jones potential) *V*_*nb*_.

Let ***r***_*i*_(*i* = 1,…, *m*) be the position vector of the m residues identified by their *C*_*α*_ Carbon atom in the reference (native) configuration, and 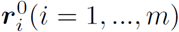 be the position vector of *m* residue in the current configuration. The peptide potential, responsible for the covalent bonds between the beads of the polymer chain, has the following expression:

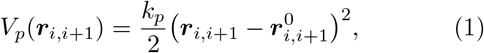

where 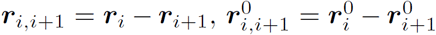 are the position vector between the bonded residues *i* and *i* +1 in the instantaneous and native configuration respectively. The norm of position vector is the bond length. The empirical constant 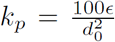 is in term of the equilibrium length parameter *d*_0_, and is in the unit of energy parameter *ϵ*. In our simulation, *d*_0_ = 3.8 Å is the average distance between two adjacent amino acids and *ϵ* sets the energy scale of the model. The bond potential is nothing more than a simple harmonic potential with spring constant *k*_*p*_.

he angular potential *V*_*θ*_ is used to recover the secondary structure of protein in reference native conformation. Mathematically, it is equivalent to peptide potential *V*_*p*_ by replacing the relative displacement by angular difference, that is

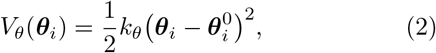

where *k*_*θ*_ = 20*ϵ* rad^-2^ is the elastic constant expressed in term of the energy computational unit *ϵ*, and *θ*_*i*_, 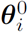 are bond angles formed by three adjacent beads in the simulated (time-dependent) and native conformation, respectively.

The dihedral potential (torsion) is 1-4 interaction, and are expressed as a function of the twisted angles *φ*_*i*_ and 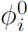, again refereed respectively to the actual and crystal configuration. The dihedral potential is important for the recovery of the correct protein secondary structure. The twisted angle is the angle formed between the two planes determined by four consecutive amino acids along the chain. The definition of twisted angle potential *V*_*φ*_ is

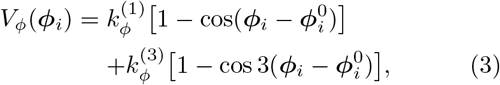

where 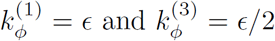 are dihedral constants expressed in term of energy unit *ϵ*.

Non-bonded (nb) interactions between nonconsecutive amino acids are modeled with Lennard-John 12–10 potential. In Gō-like model a distinction is made among the pairs of residues that interact following a potential that has also an attractive part, in addition to a repulsive part. The criteria for this distinction is made on the basis of the native distance with respect to a parameter of the model, the so-called cut-off radius, *R*_*c*_. Native contact are identified by the cut-off distance *R*_*c*_ to be chosen such that two residues *i* and *j* form a native interaction if their distance *r*_*ij*_ in the native state is less than cutoff distance *R*_*c*_. This criterion selects a certain number of native contacts on the Proteins crystallographic structure. On the other hand, when two residues are not in native contact *r*_*ij*_ > *R*_*c*_, they interact only through the Lennard-Jones repulsive tail (*σ*/*r*_*ij*_)^12^, where *σ =* 4.5 Å is a free length parameter correlated with the extension of the excluded volume (self-avoiding polymer chain). In other words, such residues in the protein chain will interact attractively if brought to a distance greater than the native one and repulsive otherwise. The expression for Lennard-Jones potential is:

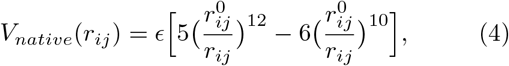

where all the symbols have been already defined. When 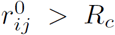, the purely repulsive contribution *V*_*nonnative*_ is assigned to the pair of amino acids considered. This term enhances the cooperatively in the folding process and takes the form of a Lennard-Jones barrier

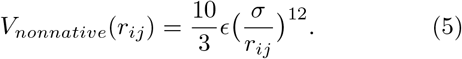

The non-bonded potential *V*_*n*__b_ summarized the possible long range interaction just described above and reads as

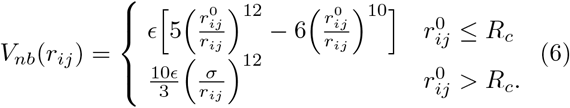

The total potential acting on all the residues of the proteins is then:

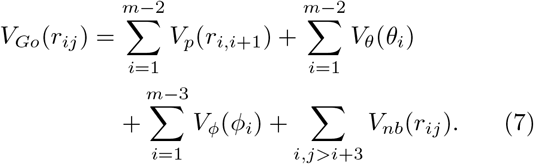

The confinement effect on protein dynamics can be represented by a step-like soft-core repulsive cylindrical potential. The cylinder axis of symmetry is set adjacent with the *x*-axis of the frame of reference used for protein trans-location simulation. The same *x*-axis direction is used to develop the mechanical pulling of the protein by using a constant force *F*_*x*_ applied to the foremost beads inside the confinement.

The expression of the pore potential is given by:

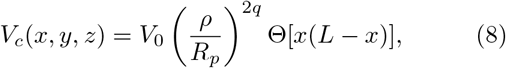

where *V*_0_ = 2*ϵ* and Θ(*s*) = [1 + tanh(*αs*)]/2 is a smooth step-like function limiting the action of the pore potential in the effective region [0, *L*]. *L* and *R*_*p*_ are pore length and radius respectively. Also, 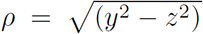 is the radial coordinate. The parameter *q* tunes the potential (soft-wall) stiffness, and *α* modulates the soft step-like profile in the *x*-direction; the larger the *α*, the steeper the step. In our simulation, we consider *q =* 1 and *α =* 2 Å^2^. The driving force *F*_*x*_ acts only in the region in front of the pore mouth x ∈ [—2,0], and inside the channel [0, *L*]. Pore length *L* = 100 Å and radius *R*_*p*_ = 10 Å are taken from αHL structure data.

**FIG. 1.**
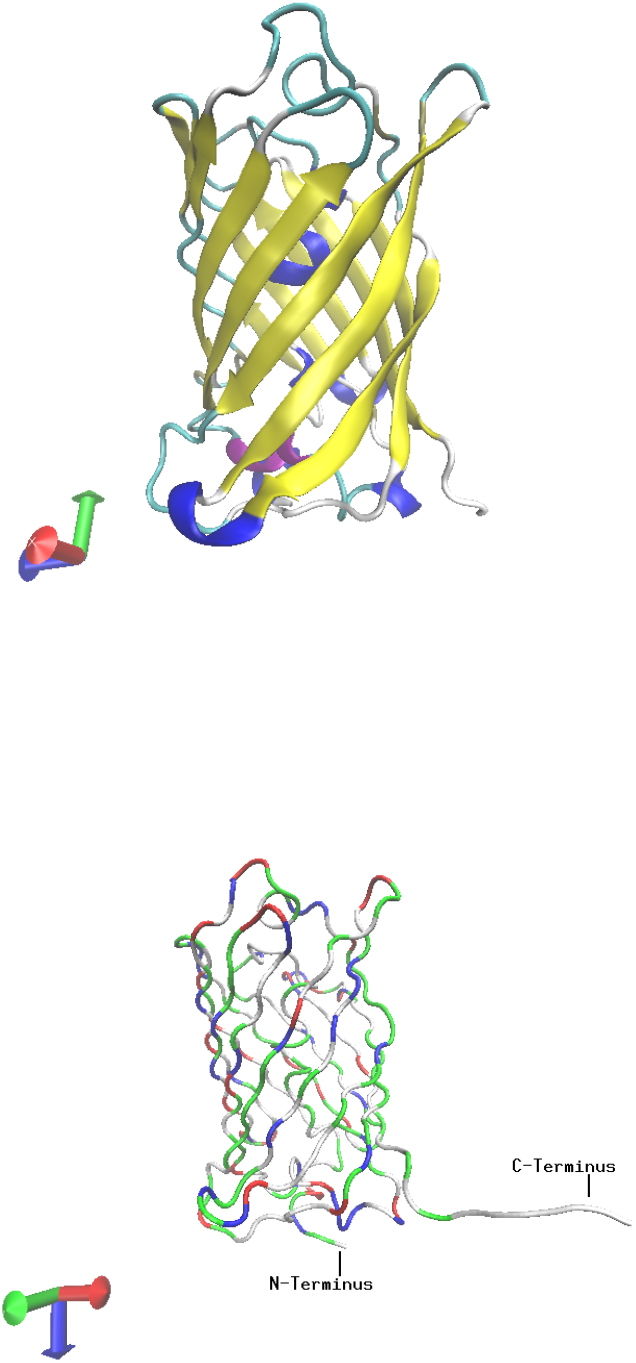
The overall shape of the GFP and its association into dimers. Eleven strands of *β*-sheet (yellow) form the walls of a cylinder. Short segments of *α*-helices (blue and purple) cap the top and bottom of the *β*-can and also provide a scaffold for the fluorophore, which is near geometric center of the can. The *β*-sheet outside and the helix inside, represent a new class of proteins. Figure produced by VMD-software. GFP with colors that vary according to the residues type. The N and C-termini are marked. To facilitate the trapping inside the confinement geometry i.e cylinder, a linker without structure are added to the C terminus. Figure produced by VMD-software.

### B. Equation of Motion: Langevin dynamics

The equation of motion which governs the motion of material point is the well-know Langevin equation which is usually employed in coarse-grained molecular dynamics simulation. Consequently, numerical investigations are performed at constant temperature. The over-damped limit is actually implemented 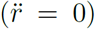 and a standard Verlet algorithm is used as numerical scheme for time integration. The Langevin equation is given by

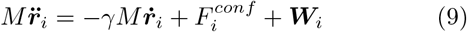

where 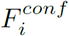 is the sum of all the internal and external forces acting on residue *i*. Here γ is the friction coefficient used to keep the temperature constant (also referred as Langevin thermostat). The random force ***W***_*i*_ accounts for thermal fluctuation, being a delta-correlated stationary and standard Gaussian process (white noise) with variance 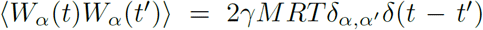. The random force satisfies the fluctuation-dissipation theory; the mean-square of *W* is proportional to the corresponding friction coefficient γ.

## III. RESULTS

### A. Unfold of Green Fluorescent Protein (GFP)

The remarkable cylindrical fold of the GFP seems ideally suited for its function to emit greeen fluorescence. The strands of the *β*-sheet are tightly fitted to each other like stave in a barrel and form a regular pattern of hydrogen bonds, as shown in Fig. (1). Together with the short *α*-helices and loops on the ends, the can-structure forms a single compact domain and does not have obvious clefts for easy access of diffusible ligands to the fluorophore. The florescence properties of GFP depend on its folded structure. Figure (2-lower panel) shows the assignment of the stages of the GFP unfolding by a constant force within the cylindrical geometry. Initially the force unravels the *β* strands 11 × 7 of GFP, generating a short pause as shown in the figure. This pause enforce the protein to enter as a loop in the cylinder. The remain *β* strands 6 × 1 are couple together and translocate as a loop.

Fig.(3) shows the unfold trajectory of GFP pulling by a constant force *F*, 0, 0 acting only on the C-terminus bead (**r**_*N*_). Defining the collective variable coordinate *Q*,

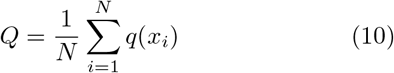

where *x*_*i*_ is the *x*-coordinate of the *i*th bead and *q*(*x*) is the piecewise function

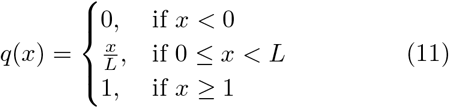

The value of *Q* = 0 corresponds to the case when the folded GFP is completely on the Cis side of the confinement geometry and *Q* = 1 when the GFP is completely unfold. The value 0 < *Q* < 1 represent the coordinate when the beads is inside the confinement effect.

Recently the experiment and computational studies of GFP have shown that braking off two adjacent *β* strands required larger force than simply unzipping them [26]. As shown in figure (2-right upper panel) the topological structure of GFP, the two *beta* stands, *β*_1_ and *β*_2_ which are parallel to each other required a larger pulling force to separate them. The remaining other *β* strands in GFP are antiparallal and the pulling force unzip them easily. As a result the GFP after unfolding up to *ß*_7_ to *ß*_1_ 1 enter as a folded loop inside the pore. The trajectory plot show a short live intermediate. This short live intermediate or pause comprising the N terminal residues that from *β* strands 1 though 6, is in good agreement with previous mechanical unfolding experiments of GFP [27].

**FIG. 2.**
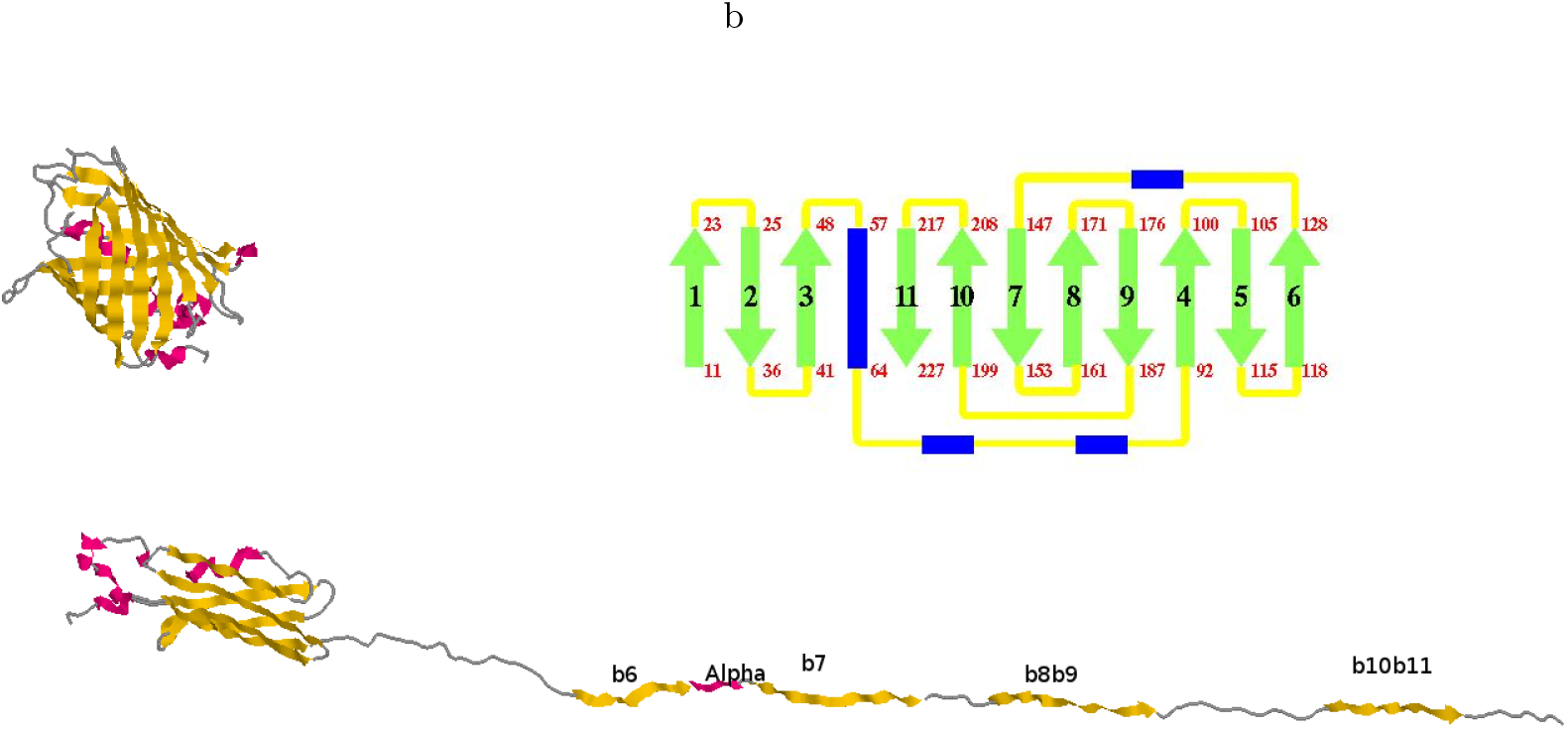
A schematic diagram of GFP unfolding and trans-locate through the *α*-hemolysin. The upper panel (left) shows the GFP molecule before unfolding. (Upper panel right) A topology diagram of the folding pattern in GFP. The *β*-sheet strands are shown in green color, α-helices in blue, and connecting loops in yellow. The position in the sequence that begin and end each major secondary structure element are also shown. The anti-parallel strands (except for the interactions between strands 1 and 6) make a tightly formed barrel. The bottom panel shows the unfolding intermediate and its trans-location through pore.

**FIG. 3.**
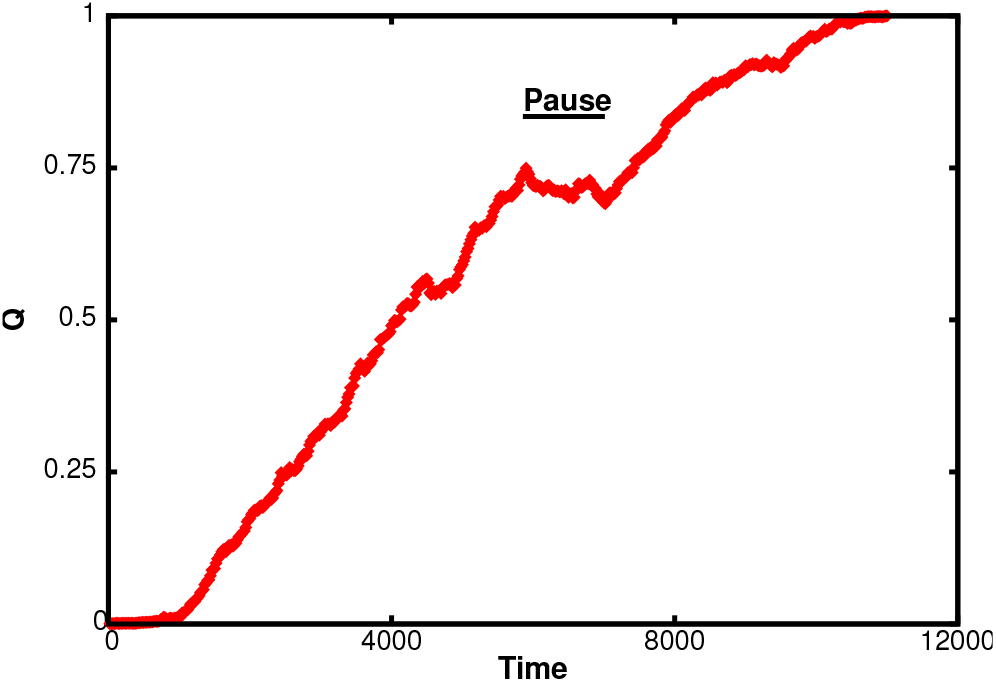
Trajectory of GFP in collective variable coordinate *Q* as a function of time during unfolding and trans-location through a static *α*-hemolysin pore effected using a constant external force of *F* = 1.5, at the C-terminus. The rip in the trajectory corresponding to a GFP unfolding is preceded by a pause or stall effect follow by a non-spontaneous unfold. After GFP is unfolded, the pore trans-locates the unfolded polypeptide chain with occasional pauses.

### B. GFP Vs MBP

Two different type of protein, one with majority of *β*-sandwich proteins (GFP) [28] and another with majority of *α*- helix proteins (Maltose Binding Protein-MBP) [29], shows significant different behavior when unfold and translocate through a channel by external force. Figure (4) shows the secondary structure, unfold and translocation of GFP (Fig.4 Panel(a)) and MBP (Fig.4 Panel(b)). We find a conformation of *β*-hairpin when translocating GFP through a static pore. This behavior can be attributed due to the anti-parallel *β*-strands at the N and C terminal. As shown in the topology structure of GFP in Fig. (2), the anti-parallel *β*-strands make a tightly formed barrel. These results are already conformed both in simulation, and in theoretical studies on different protein topologies [30, 31] that *β*-sheet protein with parallel N and C terminal strands (orthogonal to inter-strand hydrogen bonds) have greater higher mechanical stability than proteins with anti-parallel *β* strands when pulled from their N or C termini. The interpretation of this is that mechanical force applied orthogonal to inter-strand hydrogen bonds loads all bounds simultaneously before mechanical failure. On the other when force applied parallel to the hydrogen bonds leads each bond in turn, resulting a peeling rupture of each bond at relatively low force.

**FIG. 4.**
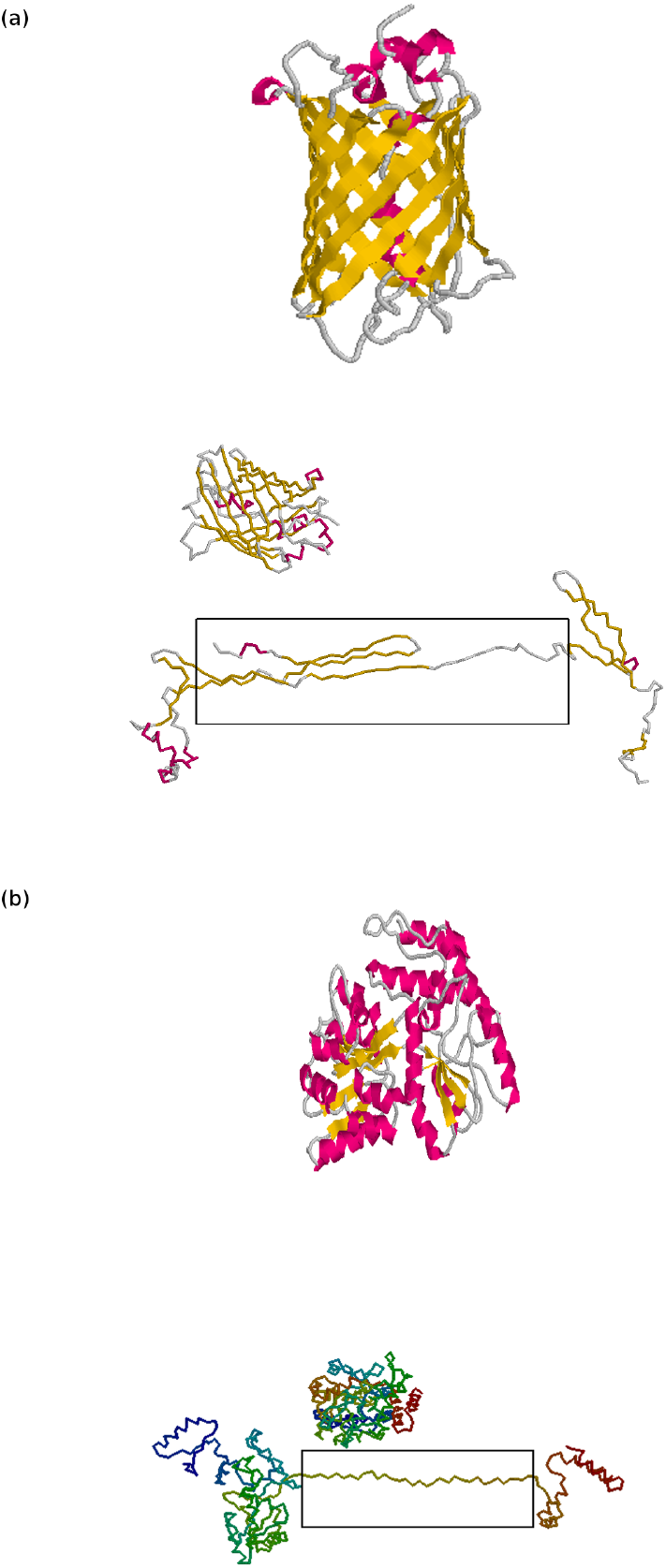
Panel (a): Structure of GFP with *β*-sheet shown in yellow color while the α-helix in pink color. The intermediate panel represent GFP coarse grained description produced in RasMol software. Lower panel (a): Schematic view of GFP translocation through a pore of length *L* = 100. Panel (b): Secondary structure of MBP with *α*-helix depicted by pink color and the *β*-sheet by yellow color. The intermediate panel describe MBP coarse grained description. Lower panel (b): Schematic representation of unfolding at the cis-side, translocation through the pore and refolding at the trans-side of MBP.

Unfolding and translocation of MBP mechanically is in contrast with the GFP molecule as shown in Fig. (4(b)). The long polypeptide chain of MBP unfold and translocation in single file conformation. The MBP unravel begins at the C-terminus where the three α-helices are mechanically complaint and unravel at very low external forces.

**FIG. 5.**
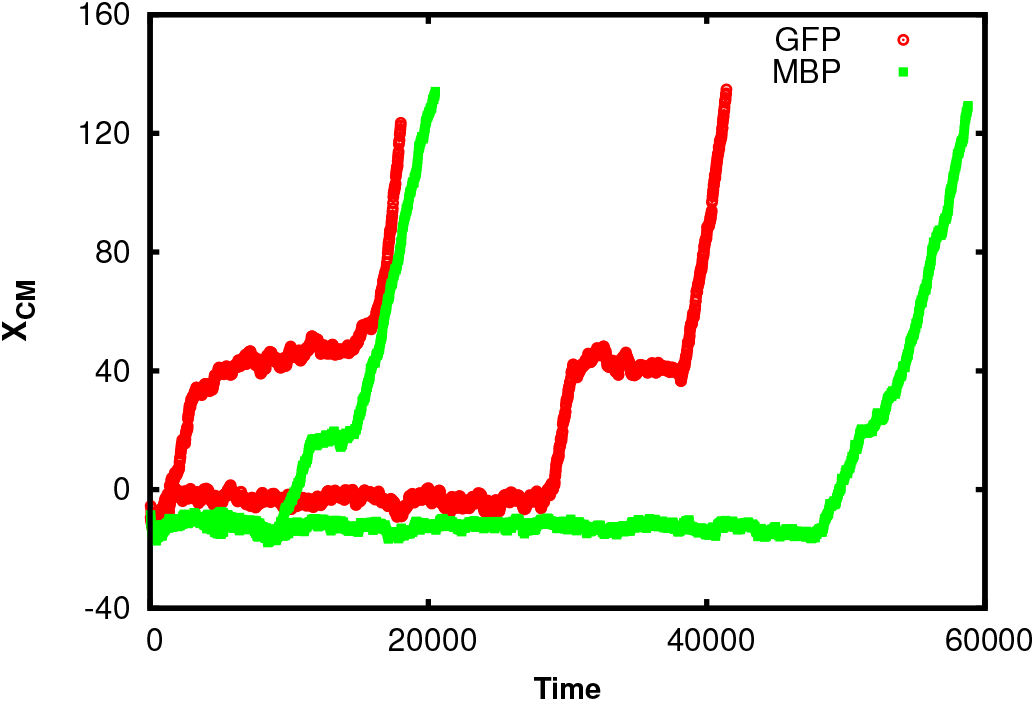
Translocation reaction coordinate as a function of time. The red empty dots represent the trajectory of GFP molecule when translocating through a static pore. The green filled dots describe the trajectory of center of mass coordinate using MBP molecule using a constant force *F* = 2.0 at the C-terminus.

On the other side, the N-terminal domain which consists of a five *β*-strands unravel at the expenses of high force (the configuration of *β*-sheets and α-helices are illustrated in Fig. (4).

Figure (5) show trajectory of center of mass coordinate (reaction coordinate *X*_*CM*_) as a function of simulation time for the GFP molecule (red empty dots), and for the MBP molecule (green filled dots) when translocating through a static pore. The rip in the trajectory corresponding to a GFP unfolding is preceded by a long pause or stall effect as compare to the short rip in the MBP trajectory.

## IV. CONCLUSION

We investigate the unfold and transport of GFP via a cylindrical geometry using coarse-grained numerical molecular dynamics simulation. Proteins with majority of *β*-sheet, such as green florescent protein, exhibit more mechanical deformation resistance as compare to *α-*helical proteins. Due to topological structure of GFP, it show a pause (stall effect) and translocate as loop configuration through nanopore. The loop configuration show significance resistance at low applied force acting along the axis of translocation from cis to trans side. On the other hand. MBP protein translocate as a single file configuration without showing any stall effect.

### Supporting Information

**Movie S1** This movie represents the translcaion of GFP through nanopore having length *L* = 100 Å and radius *R*_*p*_ = 10 Å taken from *α*HL structure data.

## ACKNOWLEDGMENTS

The author would like to thank Fabio Cecconi for helpful discussion, and Institute of Complex Systems, National Research council of Rome for providing access to computational resources.

